# Pre-Human Immunodeficiency Virus (HIV) infection Th17 CD4^+^ T cells as predictors of early HIV disease progression

**DOI:** 10.64898/2025.12.29.696811

**Authors:** Tosin E. Omole, Huong Mai Nguyen, Agata Marcinow, Naima Jahan, Giulia Severini, Nivashnee Naicker, Katherine Thomas, Connie Celum, Nelly Mugo, Andrew Mujugira, James Kublin, Lawrence Corey, Aida Sivro, Jairam Lingappa, Glenda Gray, Lyle R. McKinnon

**Author notes:** Corresponding author (TO) (LM).

## Abstract

Interleukin-17-producing T helper (Th17) CD4^+^ T cells are highly susceptible to HIV infection and are depleted early in people living with HIV. Here, we investigated whether systemic Th17 cell levels prior to HIV infection are associated with subsequent HIV disease progression. We analyzed archived cryopreserved peripheral blood mononuclear cells (PBMCs) collected within one year prior to HIV acquisition from participants enrolled in a South African cohort (HIV Vaccine Trials Network [HVTN] 503; n = 35) and an East African cohort (Partners Pre-exposure Prophylaxis/Couples’ Observational Study [PP/COS]; n = 32). Th17 cell frequencies were quantified by flow cytometry. In HVTN 503, higher pre-HIV IL-17^+^ CD4^+^ T cell frequencies were inversely correlated with CD4/CD8 ratio measured both within 180 days (Spearman rank R = -0.42, *p* = 0.012) and beyond 180 days (R = -0.55, *p* = 0.001) after HIV infection, and were associated with faster CD4^+^ T cells decline (adjusted hazard ratio [aHR] = 3.5, 95% CI: 1.2-9.9, *p* = 0.020). In contrast, no significant association with CD4 decline was observed in the PP/COS cohort (HR = 1.2, 95% CI: 0.4–3.4, *p* = 0.795). Sex-stratified analyses in HVTN 503 indicated a more pronounced association between pre-HIV IL-17^+^ CD4^+^ T cells and faster CD4 decline in males than females. In analyses combining all cohorts, higher pre-HIV IL-17^+^ CD4^+^ T cell frequencies remained associated with faster CD4 decline, particularly among younger participants (HR = 3.5; 95% CI: 1.35-9.22, *p* = 0.010). Pre-HIV IL-17^+^ CD4^+^ T cell frequencies were not associated with peak or set-point viral load in either cohort. Together, these findings suggest that pre-HIV Th17 cells abundance may influence subsequent HIV disease progression independently of early viral replication.

**Author Summary:** HIV infection leads to progressive damage of the immune system, but the rate at which this damage occurs varies widely between individuals. Understanding the factors that influence HIV disease progression is essential for improving prevention and treatment strategies. Previous studies have suggested that immune characteristics present before infection may shape disease outcomes after HIV acquisition. In this study, we examined whether the abundance of IL-17-producing CD4^+^ T cells, known as Th17 cells, measured prior to HIV acquisition, were associated with markers of disease progression after infection. We analyzed blood samples from individuals who were HIV-negative at the time of sampling and later acquired HIV during follow-up. We found that individuals with higher pre-infection levels of Th17 cells experienced faster immune decline after HIV infection, including more rapid loss of CD4^+^ T cells and lower CD4/CD8 ratios. These associations were observed independently of viral load and varied by cohort, age and sex. Our findings indicate that immune conditions present before HIV infection can influence subsequent disease progression and suggest that Th17 cells may serve as biomarkers to identify individuals at higher risk of rapid HIV-related immune damage.

## Introduction

HIV disease progression is heterogenous, and early events following HIV acquisition are important determinants of HIV reservoir formation and long-term immune parameters. The host immune profile near the time of HIV infection can shape early viral replication and the associated immune damage, which are key predictors of HIV pathogenesis [1–4]. Previous studies have shown that the abundance of HIV target cells prior to infection may accelerate disease progression [1,4,5]. Pre-HIV immune profiles characterized by high expression of the HIV-susceptible migratory marker integrin α4ꞵ7 [1,5,6], elevated immune activation markers [3] and increased plasma cytokines [4,7] have been associated with heightened HIV susceptibility and/or faster disease progression post-infection.

A subset of CD4^+^ T cells, Th17 cells, are important targets for HIV and play a crucial role in mucosal immunity, defending against bacterial and fungal infections [8–10]. Th17 cells are enriched at mucosal surfaces, where they maintain barrier integrity and produce pro-inflammatory cytokines, including interleukin-17 (IL-17). [9,10]. IL-17 mediates key host immune mechanisms such as neutrophil recruitment, regulation of mucosal epithelial permeability, and induction of antimicrobial peptides [11,12].

The gastrointestinal mucosa, a primary site for HIV replication, is highly enriched for Th17 cells [13,14]. HIV preferentially infects CD4^+^ T cells, and untreated infection is characterized by progressive CD4^+^ T cell loss, particularly in mucosal tissues [13,14]. Th17 cells are highly permissive to HIV infection and are among the first cells to be depleted during acute infection [15–17], likely due to their mucosal location and high expression of HIV coreceptors CCR5 and CXCR4 [18–20]. Early Th17 depletion contributes to mucosal barrier disruption, increased microbial translocation, and systemic immune activation, which in turn drives HIV disease progression [7,12,17,21,22]. Notably, this rapid Th17 depletion is observed in HIV-susceptible hosts such as humans and rhesus macaques, but not in natural hosts like sooty mangabeys or African green monkeys, nor in HIV-infected long-term non-progressors, who exhibit slower progression to AIDS [17].

Although Th17 cells are recognized as key targets of HIV and their depletion is linked to disease progression, it remains unclear whether the abundance of these cells in peripheral blood before infection influences subsequent HIV outcomes. In this study, we aimed to determine whether higher pre-HIV Th17 cell frequency predicts faster disease progression in two African cohorts.

## Results

### Study participants characteristics

We analyzed 67 PBMC samples from participants enrolled in the HIV Vaccine Trials Network (HVTN 503, n = 35) and from the Partners Pre-exposure Prophylaxis and Couples Observational Study cohorts (PP/COS, n = 32). All participants were HIV-seronegative at the time of blood draw but subsequently acquired HIV during longitudinal follow-up.

Participants in HVTN 503 were younger, with a median age of 23 years (Interquartile range [IQR]: 22-27) compared with those in PP/COS, who had a median age of 30 years (IQR: 25-40). The HVTN 503 cohort included 17 female and 18 male participants, while PP/COS included 19 female and 13 male participants. The median number of days from sample collection to estimated infection was 177 (IQR: 136-219) for HVTN 503 and 225 (IQR: 116-328) for PP/COS. None of the HVTN 503 participants completed the originally planned vaccination regimen, as the trial was stopped early for futility [23]. We use the acronym “PP/COS” to denote participants combined from two concurrent clinical trials involving heterosexual serodiscordant couples: Partners PrEP (n = 25) and COS (n = 7). More than 50% of PP/COS participants had a baseline risk score greater than 5, reflecting serodiscordant couples at higher risk of HIV transmission. Baseline characteristics of all participants are summarized in Table 1.

**Table 1.** Baseline characteristics of participants.

| Variables | Cohorts |  |
| --- | --- | --- |
|  | HVTN 503 (*n* = 35) | PP/COS (*n* = 32) |
| Age (years)<sup>a</sup> | 23.00 (22.00 – 27.00) | 30.00 (25.00 – 40.00) |
| Estimated sampling to infection (days)<sup>a</sup> | 177.00 (136.00 – 219.00) | 225.00 (116.00 – 328.00) |
| CD4 Count(/mm<sup>3</sup>)<sup>a</sup> | 392.50 (290.00 – 482.50) | 497.00 (428.50 – 638.50) |
| Sex |  |  |
| Female | 17 (48.57) | 19 (59.38) |
| Male | 18 (51.43) | 13 (40.62) |
| Treatment Arm |  |  |
| Placebo | 17 (48.57) | 14 (43.75) |
| Vaccine | 18 (51.43) | N/A |
| PrEP | N/A | 11 (34.38) |
| HSV-2 Status |  |  |
| Negative | 15 (42.86) | 4 (12.50) |
| Positive | 20 (57.14) | 17 (53.13) |
| Circumcision (Male) | 6 (17.14) | 0 (0.00) |
| Treatment Status (Off treatment early) | 35 (100.00) | N/A |
| Vaccine Dose |  |  |
| 1 | 7 (20.00) | N/A |
| 2 | 28 (80.00) | N/A |
| AD5 Titre at Baseline |  |  |
| ≤ 200 | 12 34.29) | N/A |
| > 200 | 23 (65.71) | N/A |
| **Baseline Risk score** |  |  |
| < 5 | N/A | 13 (40.62) |
| ≥ 5 | N/A | 19 (59.38) |
| **Sites: South Africa** |  |  |
| Cape town | 9 (25.71) | N/A |
| eThekwini | 7 (20.00) | N/A |
| Klerksdorp | 6 (17.14) | N/A |
| Medunsa | 4 (11.43) | N/A |
| Soweto | 9 (25.71) | N/A |
| PHRU | N/A | 2 (6.25) |
| **Sites: Uganda** |  |  |
| Jinja | N/A | 6 (18.75) |
| Kampala | N/A | 13 (40.63) |
| **Sites: Kenya** |  |  |
| Kisumu | N/A | 4 (12.50) |
| Thika | N/A | 2 (6.25) |
| Nairobi | N/A | 5 (15.62) |

Data are presented as sample sizes (n) and corresponding column percentages (%) unless otherwise stated.
\*Data are presented as median (Interquartile Range).
Abbreviations: HVTN = HIV Vaccine Trials Network; PP/COS = Partners PrEP/Couples Observational Study; PrEP = Pre-exposure Prophylaxis; HSV-2 = Herpes Simplex Virus-2; AD5 = Adenovirus Type 5; PHRU = Perinatal HIV Research Unit (Soweto, South Africa); N/A = Not Applicable.

### Pre-HIV IL-17^+^ CD4^+^ T cells predicts faster CD4^+^ T cell decline after HIV infection

We examined the association between pre-HIV Th17 cells, defined as IL-17-expressing CD4^+^ T cells (Fig 1A, S1 Fig) and CD4^+^ T cell decline, a marker of HIV disease progression. In HVTN 503 cohort, higher pre-HIV IL-17^+^ CD4^+^ T cell frequencies were significantly associated with faster CD4 decline. Specifically, individuals with pre-HIV IL-17^+^ CD4^+^ T cell frequencies above the median experienced nearly a three-fold faster rate of CD4 decline in an unadjusted model (hazard ratio [HR] = 2.9, 95% confidence interval [CI]: 1.2–6.9, *p* = 0.015; Fig 1B). The association remained significant after adjustment for peak viral load (adjusted HR [aHR] = 2.5, 95% CI: 1.1–6.1, *p* = 0.038; S1 Table). In a fully adjusted model that included sex, adenovirus type 5 (Ad5) titer, herpes simplex virus-2 (HSV-2) serostatus, age, and peak viral load, pre-HIV IL-17^+^ CD4^+^ T cells continued to predict faster CD4 decline (aHR = 3.5, 95% CI: 1.2–9.9, *p* = 0.020; Table 2). Given that viral load may mediate the association between Th17 cells and HIV disease progression, we also evaluated a model excluding peak viral load. In this model, pre-HIV IL-17^+^ CD4^+^ T cell frequency remained a significant predictor of faster CD4 decline (aHR = 4.3, 95% CI: 1.6–11.9, *p* = 0.005, S2 Table).

**Figure 1.**
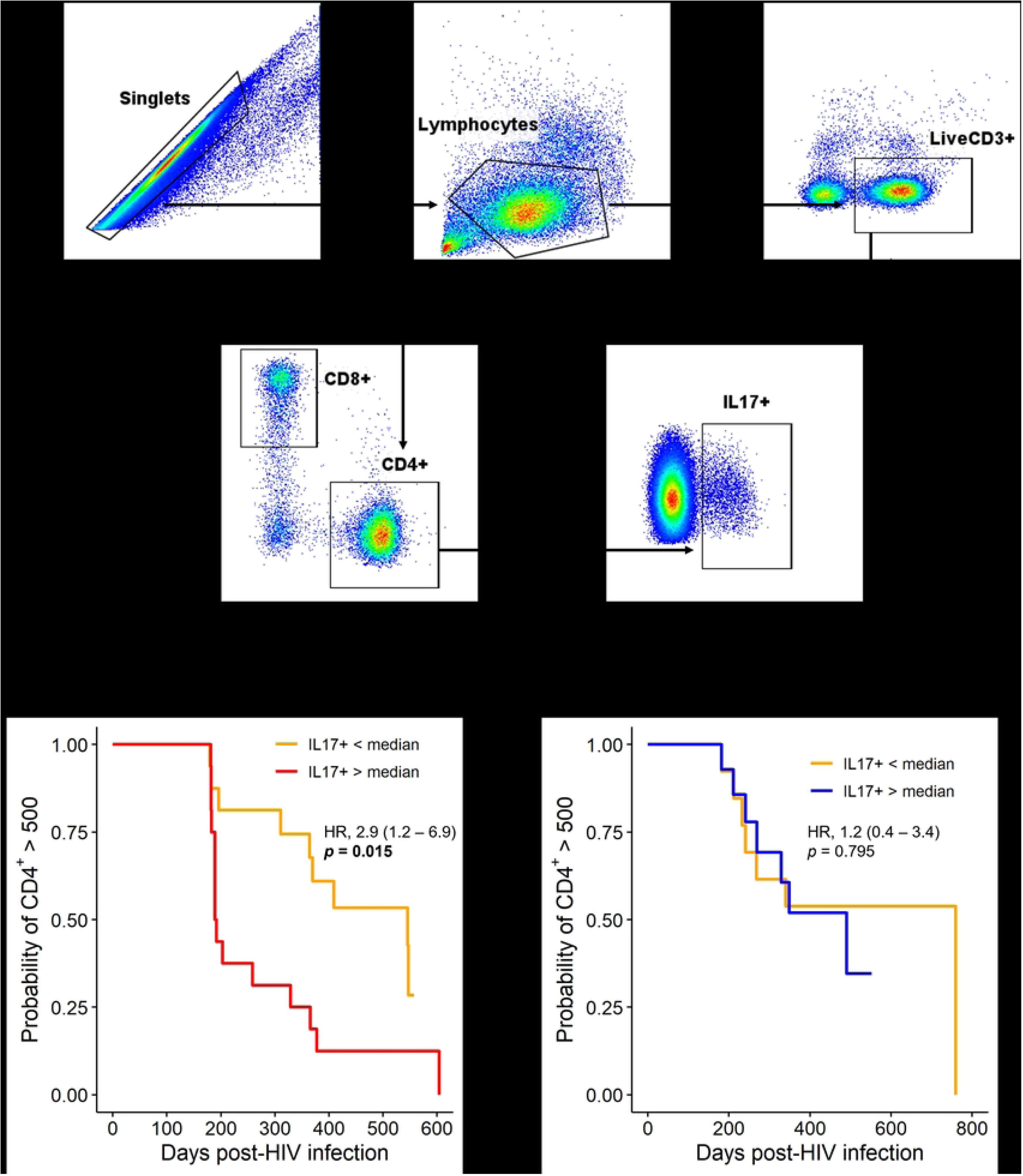
Association between pre-HIV Th17 cells and CD4^+^ T cell decline below 500 cells/mm^3^. (A) Representative flow cytometry gating strategy for IL-17^+^ CD4^+^ T cells (B) Association between pre-HIV IL-17^+^ CD4^+^ T cell frequency and CD4^+^ T cell decline in HVTN 503 (n = 32) (C) Association between pre-HIV IL-17^+^ CD4^+^ T cell frequency and CD4^+^ T cell decline in PP/COS (n = 27). CD4 counts measured after ART initiation, after 1-year post-infection, or within the first 180 days post-infection were excluded. Hazard ratios (HR) were estimated using unadjusted Cox proportional hazards model. Two-tailed *p* values are shown; statistical significance was defined as *p* < 0.05.

**Table 2.** Association between pre-HIV IL-17^+^ CD4^+^ T cells and CD4 decline below 500 cells/mm^3^.

| Cohort | HR (95% CI) | *p* value | aHR (95% CI) | *p* value |
| --- | --- | --- | --- | --- |
| **HVTN 503** | 2.9 (1.2 – 6.9) | **0.015** | 3.47 (1.21 – 9.95) | **0.020** |
| **PP/COS** | 1.2 (0.39 – 3.4) | 0.795 | 1.14 (0.32 – 4.09) | 0.844 |
| **Combined** | 2.2 (1.1 – 4.3) | **0.023** | 2.2 (0.98 – 5.01) | 0.057 |

Hazard ratios were estimated using Cox proportional hazards model. Multivariable models for HVTN 503 were adjusted for sex, adenovirus type 5 (Ad5) titer, herpes simplex virus-2 (HSV-2) status, age, and peak viral load. Models for PP/COS and the combined cohorts were adjusted for age, sex, HSV-2 status, and peak viral load. Two-tailed *p* values are shown; statistical significance was defined as *p* < 0.05. Abbreviations: HR = Hazard Ratio; aHR = Adjusted Hazard Ratio; CI = Confidence Interval; HVTN = HIV Vaccine Trials Network; PP/COS = Partners PrEP/Couples Observational Study.

In contrast, no association was observed between pre-HIV IL-17^+^ CD4^+^ T cells and CD4 decline in PP/COS (HR = 1.2, 95% CI: 0.4–3.4, *p* = 0.795; Fig 1C). When data from both cohorts were combined, higher pre-HIV IL-17^+^ CD4^+^ T cell frequencies were associated with faster CD4 decline in the unadjusted model (HR = 2.2, 95% CI: 1.1–4.3, *p* = 0.023) and showed a statistical trend in the adjusted model (aHR = 2.2, 95% CI: 0.98–5.0, *p* = 0.057; Table 2).

To assess whether this association extended to other CD4^+^ cytokine-producing subsets, we examined TNF-α, IFN-γ, GM-CSF, and IL-22 expression gated directly on total CD4^+^ T cells (S2 Fig). None of these subsets were associated with CD4 decline, indicating that the observed effect was specific to IL-17^+^ CD4^+^ T cells (S3 Table).

### Pre-HIV IL17^+^ CD4^+^ T cells inversely correlate with CD4/CD8 ratio after HIV infection

Next, we examined the relationship between pre-HIV IL-17^+^ CD4^+^ T cell frequencies and post-HIV CD4/CD8 ratios in the HVTN 503 cohort. Mean CD4/CD8 ratios were calculated separately for the first 180 days following infection and for periods beyond 180 days, and associations were also assessed using the CD4/CD8 ratio measured at the last available pre-ART visit (Fig 2). We observed a significant inverse correlation between pre-HIV IL-17^+^ CD4^+^ T cell abundance and mean CD4/CD8 ratio within the first 180 days post-infection (R = -0.42, *p* = 0.012; Fig 2A). This association became stronger beyond 180 days post-infection (R = - 0.55, *p* = 0.001; Fig 2B). This observation also holds true for CD4/CD8 ratio measured at the last available sampling visit (Figs 2C and 2D).

**Figure 2.**
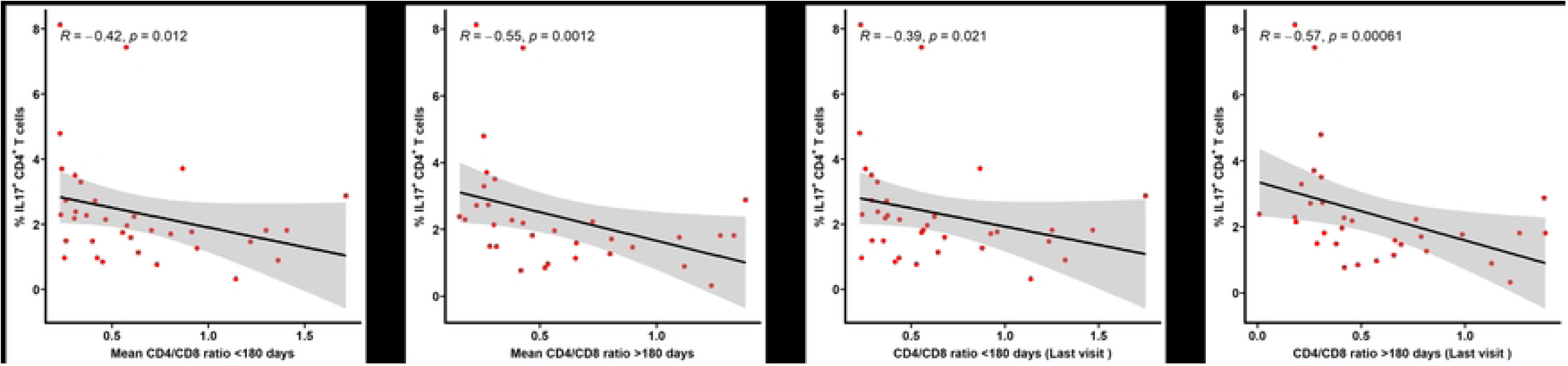
Correlation between pre-HIV Th17 cells and CD4/CD8 ratio in HVTN 503. (A, B) Correlations with mean CD4/CD8 ratio. (C, D) Correlations with CD4/CD8 ratio at last available sampling visit. CD4/CD8 ratios were calculated from absolute CD4 and CD8 counts measured within (n = 35) or after the initial (n = 32) 180 days post-infection, as indicated. Measurements obtained after ART initiation or beyond 1-year post-infection were excluded. Correlations were assessed using Spearman rank correlation (R), with linear regression lines shown for visualization. Two-tailed *p* values are shown; statistical significance was defined as *p* < 0.05.

Within the first 180 days after infection, multiple IL-17^+^ CD4^+^ T cell subsets including IL-17^+^ IFNγ^+^, IL-17^+^ IFNγ^-^, IL-17^+^ IL-22^-^ and IL-17^+^ GM-CSF^+^ were significantly correlated with mean CD4/CD8 ratio (S3 Fig). Beyond 180 days post-infection, most measured IL-17^+^ CD4^+^ T cell subsets remained significantly associated with mean CD4/CD8 ratio (S4 Fig).

In contrast to our findings for CD4 decline, IL-22^+^, IFN-γ^+^ and GM-CSF^+^ CD4^+^ T cell subsets were also inversely correlated with CD4/CD8 ratio (S5 Fig).

### Interaction and stratified analyses

To determine whether these associations extended to our entire study population, we evaluated interactions between pre-HIV IL-17^+^ CD4^+^ T cell frequencies and selected variables, including age, sex, and viral load. No significant interactions were observed when cohorts were analyzed separately. However, when both cohorts were combined, we detected a significant interaction between IL-17^+^ CD4^+^ T cells and age (*p* = 0.022). To further characterize this interaction, participants were stratified by age using the cohort median (<26 vs. ≥26 years). Among participants younger than 26 years, higher pre-HIV IL-17^+^ CD4^+^ T cell levels were associated with faster CD4^+^ T cell decline (HR = 3.5; 95% CI: 1.35–9.22, *p* = 0.010; Table 3). In contrast, no significant association was observed among participants older than the median age (HR = 1.3; 95% CI: 0.48–3.49, *p* = 0.615; Table 3).

**Table 3.** Association between pre-HIV IL-17^+^ CD4^+^ T cells and CD4 decline below 500 cells/mm^3^, stratified by age (combined cohorts)

| Cohort | < Median Age | | ≥ Median Age | |
| --- | --- | --- | --- | --- |
| | HR (95% CI) | $p$ value | HR (95% CI) | $p$ value |
| Combined | 3.5 (1.35 – 9.22) | **0.010** | 1.3 (0.48 – 3.49) | 0.615 |

Sample size for age strata were: < median age (n = 29) and ≥ Median age (n = 30), with the median age = 26 years. Hazard ratios were estimated using unadjusted Cox proportional hazards model. Two-tailed $p$ values are shown; statistical significance was defined as $p < 0.05$ . Abbreviations: HR = Hazard Ratio; aHR = Adjusted Hazard Ratio; CI = Confidence Interval.

In HVTN 503, but not in the combined cohort analyses, sex-stratified models indicated that the association between pre-HIV IL-17^+^ CD4^+^ T cells and faster CD4 decline was more pronounced in male participants (HR = 7.0, 95% CI: 1.46–33.7, *p* = 0.015; Table 4) compared with female participants (HR = 1.8, 95% CI: 0.454–7.36, *p* = 0.396; Table 4). Consistent with these findings, a significant inverse correlation between pre-HIV IL-17^+^ CD4^+^ T cell phenotypes and CD4/CD8 ratio was observed in male participants within HVTN 503, but not in female participants (Fig 3A, S6 and S7 Figs). Taken together, these findings suggest sex-and age-dependent differences in the association between elevated pre-HIV Th17 cell levels and markers of HIV disease progression, particularly within the HVTN 503 cohort.

**Figure 3.**
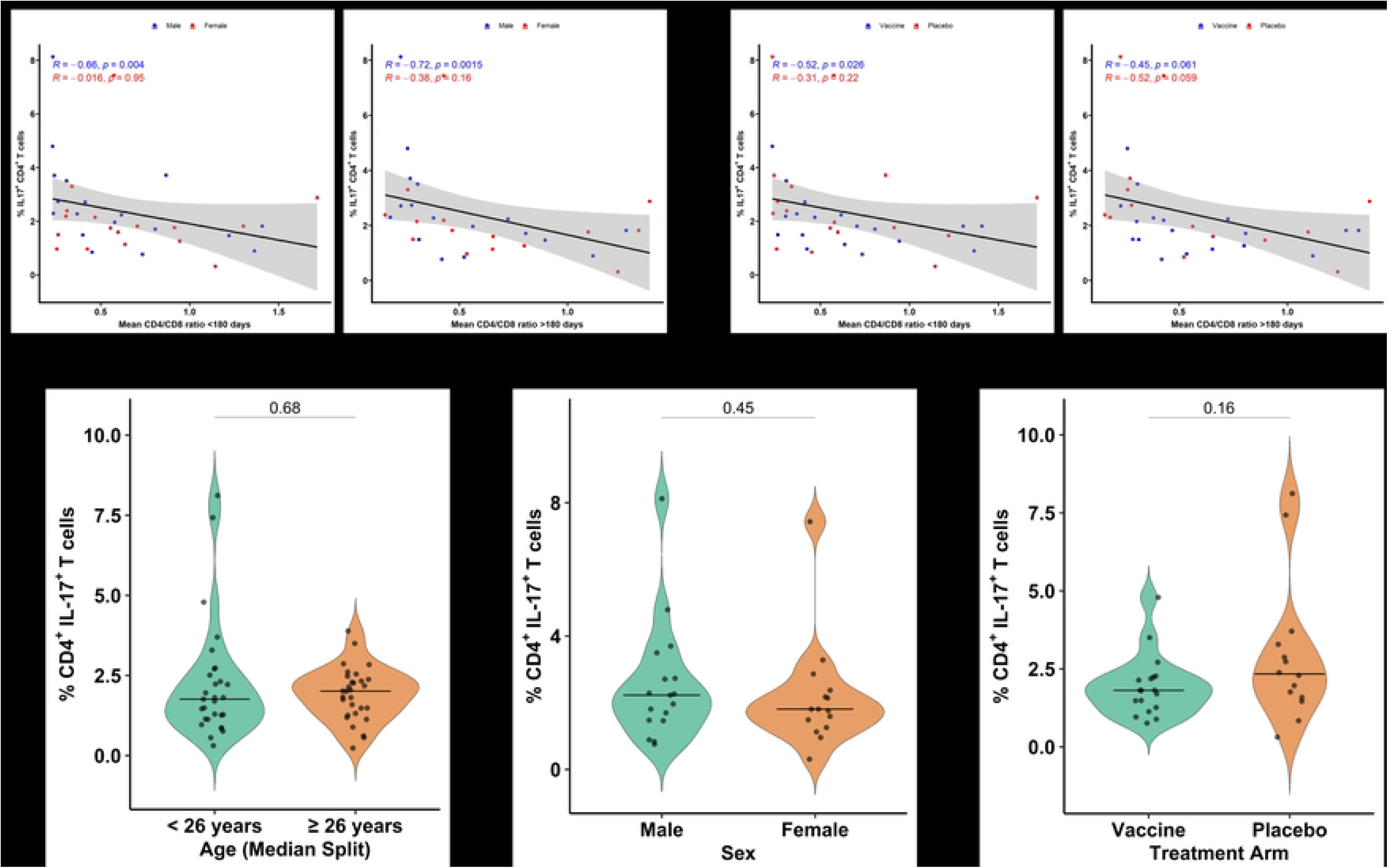
Stratified analyses of pre-HIV Th17 cells and CD4/CD8 ratio. (A) Sex-stratified correlations between pre-HIV IL-17^+^ CD4^+^ T cell frequencies and mean CD4/CD8 ratio in HVTN 503. (B) Treatment-arm-stratified correlations between pre-HIV IL-17^+^ CD4^+^ T cell frequencies and mean CD4/CD8 ratio in HVTN 503. (C) Comparison of pre-HIV IL-17^+^ CD4^+^ T cell frequencies stratified by age group (<26 vs ≥26 years) in the combined cohort. (D) Comparison of pre-HIV IL-17^+^ CD4^+^ T cell frequencies stratified by sex in HVTN 503. (E) Comparison of pre-HIV IL-17^+^ CD4^+^ T cell frequencies stratified by treatment arm in HVTN 503. CD4/CD8 ratios were calculated from absolute CD4 and CD8 counts measured within (n = 35) or after the initial (n = 32) 180 days post-infection, as indicated. Measurements obtained after ART initiation or beyond 1-year post-infection were excluded. Correlations were assessed using Spearman rank correlation (R), with linear regression lines shown for visualization. Group comparisons were performed using Mann-Whitney test. Two-tailed *p* values are shown; statistical significance was defined as *p* < 0.05.

**Table 4.** Association between pre-HIV IL-17^+^ CD4^+^ T cells and CD4 decline below 500 cells/mm^3^, stratified by sex.

| Cohort | Male Strata |  | Female Strata |  |
| --- | --- | --- | --- | --- |
|  | HR (95% CI) | *p* value | HR (95% CI) | *p* value |
| **HVTN 503** | 7.00 (1.46 – 33.70) | **0.015** | 1.8 (0.45 – 7.36) | 0.396 |
| **PP/COS** | 0.91 (0.13 – 6.47) | 0.924 | 1.6 (0.42 – 5.82) | 0.510 |
| **Combined** | 2.2 (0.80 – 5.8) | 0.127 | 2.2 (0.84 – 5.85) | 0.107 |

Sample size for sex strata were male (*n* = 17) and female (*n* = 15). Hazard ratios were estimated using unadjusted Cox proportional hazards model. Two-tailed *p* values are shown; statistical significance was defined as *p* < 0.05. Abbreviations: HR = Hazard Ratio; aHR = Adjusted Hazard Ratio; CI = Confidence Interval; HVTN = HIV Vaccine Trials Network; PP/COS = Partners PrEP/Couples Observational Study.

Next, we examined whether the associations between pre-HIV IL-17^+^ CD4^+^ T cell subsets and CD4/CD8 ratio differed by treatment arm in HVTN 503. Stratified analyses revealed heterogenous associations, with some IL-17^+^ CD4^+^ T cell subsets showing significant correlations with CD4/CD8 ratio in the vaccine arm, others in the placebo arm, and several subsets showing no significant associations in either arm (Fig 3B, S8 and S9 Figs). Hence, no consistent or uniform pattern of association across treatment arms was observed.

To determine whether differences in association across subgroups reflected underlying differences in IL-17^+^ CD4^+^ T cell expression, we compared pre-HIV IL-17^+^ CD4^+^ T cell frequencies between age (<26 vs. ≥26 years), sex, and treatment-arm subgroups. No significant differences in IL-17^+^ CD4^+^ T cell expression were observed across these subgroups (Figs 3C–3E). These findings suggest that the observed subgroup-specific associations are unlikely to be explained by baseline differences in IL-17^+^ CD4^+^ T cell expression.

### Pre-HIV IL17^+^ CD4^+^ T cells are not associated with viral load (VL)

We next assessed the correlation between pre-HIV IL-17^+^ CD4^+^ T cell frequency and viral load measures, including peak and set point viral loads. Pre-HIV IL-17^+^ CD4^+^ T cell frequencies were not significantly associated with either peak or set point viral load in either cohort (Fig 4, S10 and S11 Figs). These findings suggest that the observed association between pre-HIV IL-17^+^ CD4^+^ T cells and disease progression is unlikely to be mediated by differences in early viral replication.

**Figure 4.**
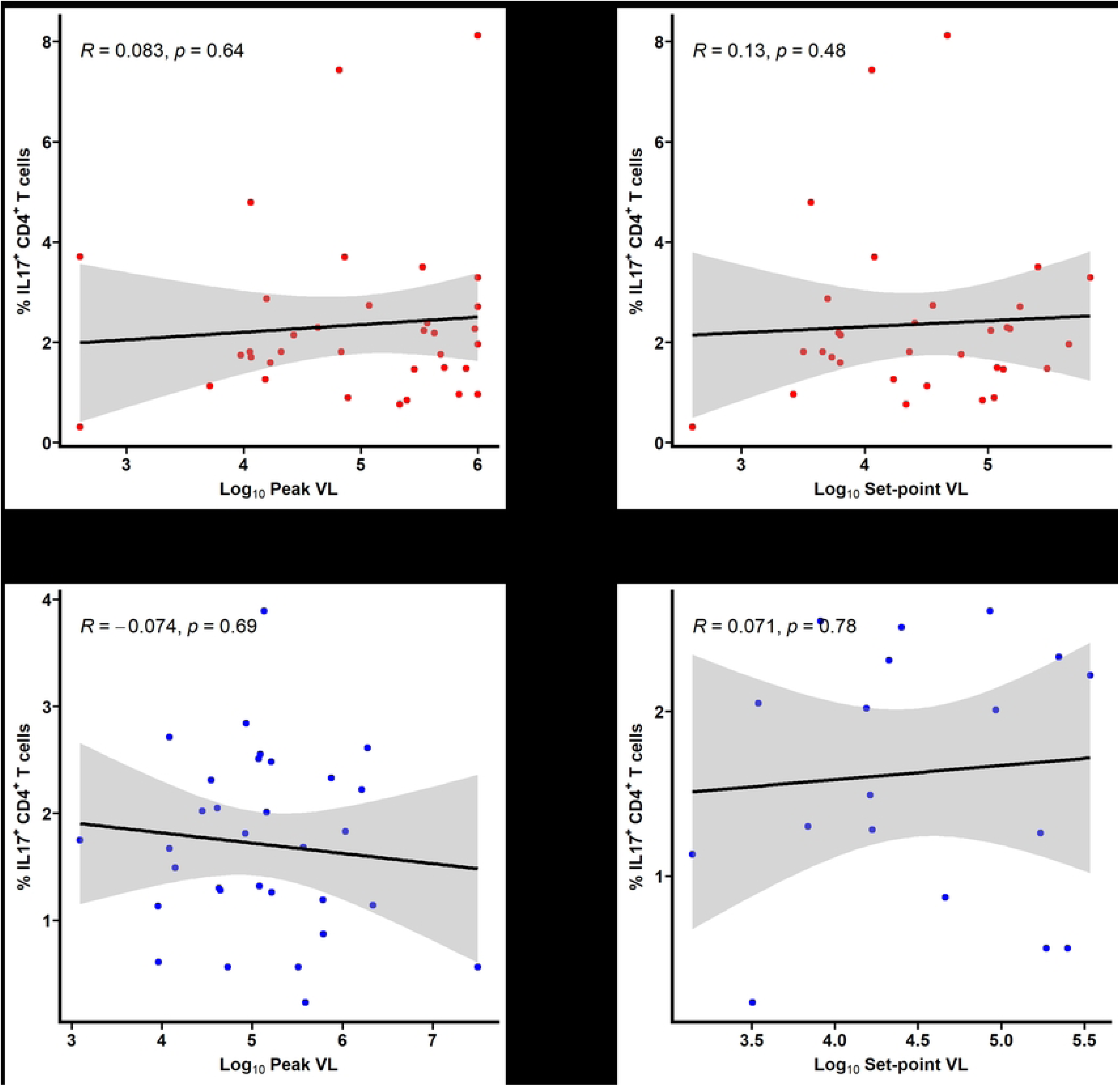
Correlation between pre-HIV Th17 cells and viral load. **(A, B)** Correlation with peak (n = 35) and set point (n = 32) viral loads in HVTN 503. **(C, D)** Correlation with peak (n = 32) and set-point (n = 18) viral loads in PP/COS. Peak viral load was defined as the highest viral load measured within the first 180 days post-infection, while set-point viral load as the mean viral load after 180 days post-infection. Viral load measurements obtained after ART initiation or beyond 1-year post-infection were excluded. Correlations were assessed using Spearman rank correlation (R), with linear regression lines shown for visualization. Two-tailed *p* values are shown; statistical significance was defined as *p* < 0.05.

## Discussion

Our study demonstrates that higher pre-HIV circulating IL-17^+^ CD4^+^ T cells (Th17 cells) predict faster CD4^+^ T cell decline and lower CD4/CD8 ratios, particularly in the HVTN 503 cohort. Notably, pre-HIV Th17 frequency was not associated with peak or set-point viral loads, indicating that being predictive of disease progression is not mediated solely by viremia. These findings suggests that the pool of susceptible Th17 cells present prior to infection may accelerate early immune damage independently of viral load. This observation is consistent with prior studies showing that baseline immune parameters, particularly the abundance of HIV target cells, can influence subsequent disease progression [1,4,5,7,24]. In contrast to previous findings that reported higher pre-SIV Th17 cells being correlated to lower peak and set-point viral loads [2], our results suggest that the effect of Th17 cells on disease progression is largely independent of HIV viremia and may instead reflect their role in promoting immune activation and systemic inflammation.

Systemic inflammation and immune activation are major drivers of immune dysfunction in HIV, contributing to rapid disease progression in the absence of treatment [7,17,25]. Given Th17 cells are highly permissive to HIV infection, a higher pre-infection frequency of Th17 cells may therefore expand the pool of target cells available to the virus, accelerating early CD4^+^ T cell loss [17,19,25,26]. Given their role in maintaining mucosal barrier integrity and regulating immune responses to bacterial pathogens [8,13], early Th17 depletion may exacerbate microbial translocation and systemic immune activation, further driving CD4^+^ T cell decline and disease progression.

Stratified analyses revealed age- and sex-specific differences in these associations. Younger participants (<26 years) showed stronger associations between pre-HIV Th17 cell frequency and CD4 decline when cohorts were combined, suggesting potential age-dependent immune susceptibility. However, this age-specific effect was not observed when cohorts were analyzed separately and may partly reflect differences in cohort composition, as the PP/COS cohort comprised an older population and did not show a significant association. Therefore, this observation should be interpreted cautiously and requires confirmation in larger, more age-balanced cohorts.

In HVTN 503, male participants exhibited more pronounced associations between pre-HIV Th17 cell frequency and both accelerated CD4 decline and lower CD4/CD8 ratios. Sex-based differences in HIV disease outcomes have been previously reported, with men showing lower baseline CD4 counts and distinct early immune responses compared to women during acute infection [27–30]. Our findings in pre-HIV Th17 cell biology may contribute to these divergent disease outcomes. Importantly, sex bias in the Th17 pathway is well documented across multiple disease contexts [31–33]. Experimental studies have shown that androgen receptor signaling can modulate Th17 differentiation and IL-17 production through metabolic and transcriptional regulation, leading to sex-dependent differences in Th17 effector function [34,35]. Conversely, females often exhibit enhanced Th17-mediated inflammation and have a higher prevalence of Th17-driven autoimmune diseases, suggesting a sex-based regulation of this pathway. These differences have been attributed to estrogen-mediated promotion of Th17 differentiation and reduced androgen-mediated restraint [34,36–38]. Such hormonally mediated differences in Th17 cell function may contribute to divergent immune set points between men and women and provide a biological explanation for the sex-specific associations observed in our study. Further investigations are warranted to determine how sex-dependent regulation of Th17 cells influences early HIV pathogenesis and disease progression.

While our findings emphasize a role for circulating Th17 cells in HIV disease, several limitations warrant consideration. We focused on circulating Th17 cells, which may not fully reflect immune dynamics at mucosal sites where HIV replication predominantly occurs. Our study may also be underpowered to detect interaction effects within individual cohorts, and confirmation of these findings will require larger studies. Missing CD8 data in one cohort limited comprehensive CD4/CD8 ratio analyses, and the geographic focus on African cohorts may limit generalizability. We were also unable to study post-HIV infection immune parameters apart from CD4 and CD8 proportions, limiting our ability to understand how Th17 cells may predict additional immune features of HIV progression including HIV-specific responses and immune activation.

## Conclusion

In conclusion, elevated pre-HIV IL-17^+^ CD4^+^ T cell levels were associated with accelerated HIV disease progression, independent of viral load, with cohort-, age- and sex-specific effects. These findings suggest that pre-infection Th17 cell abundance may serve as a biomarker of early HIV progression risk. Future studies are needed to elucidate the mechanisms underlying these associations and to explore whether targeting Th17-related pathways could potentially modulate HIV disease outcomes.

## Methods

### Study cohorts

Study participants were enrolled from two independent cohorts situated in South and East Africa, namely, HVTN 503 (N = 35) and Partners PrEP (N = 25), respectively. A subset of samples from Couples Observational Study (N = 7), another East African cohort similar to Partners PrEP, was included in our analyses. Hence, the renaming of our East African cohorts to PP/COS (N = 32). Details about each cohort and participants enrolment have been described elsewhere [23,39–42]. Study cohorts comprise of heterosexual, high-risk, HIV negative men and women above 18 years from multiple sites in South Africa (Cape Town, Soweto, MEDUNSA, eThekwini and Klerksdorp-Orkney-Stilfontein-Hartbeesfontein [KOSH]) for HVTN 503 and Kenya (Kisumu, Nairobi, Thika, Eldoret) and Uganda (Kampala, Jinja, Kabwohe, Mbale, Tororo) for PP/COS. Specifically, PP/COS comprised of couples in a serodiscordant relationship. Peripheral blood mononuclear cells (PBMCs) collected prior to HIV infection and cryopreserved were analyzed from participants who seroconverted during the original study follow-up.

### Sample processing

Cryopreserved PBMCs were thawed, washed and rested in warm media (RPMI 1640 supplemented with 10% fetal bovine serum and 1% Penicillin-Streptomycin) for 3 hours at 37^0^C and 5% CO_2_. Cell count and viability estimate were done manually using a hematocytometer. Cells were stimulated for 4 hours at 37^0^C with phorbol 12-myristate 13-acetate (PMA; 100 ng/ml) and ionomycin (1 μg/ml). Protein secretion was inhibited by the addition of pre-titrated amount of BD GolgiStop (Monensin) and GolgiPlug (Brefeldin A) during stimulation.

### Intracellular cytokine staining and flow cytometry analyses

Following stimulation, cells were first stained with pre-titrated amount of LIVE/DEAD^TM^ Fixable Aqua Dead Cell Stain Kit (Thermofisher Scientific) for 20 minutes to discriminate between live and dead cells. This was then followed by surface staining with a premixed cocktail of extracellular fluorochrome-conjugated monoclonal antibodies, including CD3-APCH7, CD4-BUV496, CD8-BV650, CD45RO-BUV395 and Integrin ꞵ7-BUV737, for 20 minutes at 4^0^C. Stained cells were washed, permeabilized using BD Cytofix/Cytoperm at 4^0^C for 20 minutes and stained with a premixed cocktail of intracellular cytokine antibodies, including IL-17A-BV421, IL-22-PE, GM-CSF-PECF594, IFN-γ-PECY7, TNF-α-FITC, IL-4-BV786 and IL-10-BV711, at 4^0^C for 30 minutes. Detailed information on the flow cytometry panel of antibodies used for immunophenotyping are shown in S4 Table. Cells were then washed and acquired on a BD LSRFortessa^TM^ Flow Cytometer. Immunophenotyping analysis was performed on FlowJo^TM^ version 10.6.1 (BD Life Sciences). All experiments were performed in a blinded fashion.

### Th17 cell characterization

Th17 cells were defined by flow cytometry as IL-17^+^ CD4^+^ T cells. Additional cytokines produced by these cells, including IL-22, TNF- α, IFN- γ, and GM-CSF, were also measured, and corresponding subsets were gated in relation to IL-17A expression.

### Statistical analyses

Demographic data were summarized as median (interquartile ranges) and percentages for continuous and categorical variables, respectively. Pre-HIV infection samples collected within 1 year before the evidence of HIV infection were used to analyze the correlation immune subsets and various parameter of HIV progression, including peak and set-point VL, CD4/CD8 ratio within and beyond the initial 180 days post-infection and the decline in CD4^+^ T cell counts measured post-infection, before the initiation of ART. Spearman rank correlation test was used to evaluate the correlation with HIV peak and set-point viral loads. Peak VL was defined at the highest VL within the first 180 days post-infection prior to ART initiation, while set-point VL was defined as the average VL more than 180 days post-infection before ART initiation. For HVTN 503 only, Spearman correlation test was used to evaluate the correlation between IL-17-expressing CD4^+^ T cell frequencies and mean CD4/CD8 ratio and CD4/CD8 ratio at the last available sampling visit post-HIV infection before ART initiation. Cox regression analysis was used to predict time to CD4^+^ T cell decline, with the end point defined as CD4^+^ T cell counts below 500 copies/ mm^3^ before ART initiation. The main exposure variable, pre-HIV IL-17^+^ CD4^+^ T cells frequency was dichotomized around the median expression level. CD4^+^ T cell counts measured during the first 180 days of infection were censored to exclude the transient CD4 count changes characteristic of early HIV infection [1]. CD4 counts measured after ART initiation were also excluded. All statistical analyses and plots were computed in RStudio 4.2.2. All statistics are two-tailed and *p* values ≤0.05 were considered statistically significant.

## Study approval

All participants in the parent cohorts provided informed written consent to have specimens stored for future immunological research, and the sub-studies reported here were approved by the institutional review board (IRB) at the University of Manitoba and at local IRBs where the study was conducted, where required.

## Data availability

Data will be made available upon reasonable request to the corresponding author.

## Authors contributions

Study design and funding acquisition were developed by Aida Sivro, Glenda Gray, Jairam Lingappa, Lyle R. McKinnon. Data acquisition and flow cytometry analysis were performed by Tosin E. Omole, Huong Mai Nguyen, Agata Marcinow. Data analysis and interpretation were the responsibility of Tosin E. Omole, Aida Sivro, Katherine Thomas, Lyle R. McKinnon. Study logistics and provision of reagents were coordinated by Naima Jahan, Giulia Severini. HVTN 503 cohort was managed by James Kublin, Lawrence Corey, Nivashnee Naicker, Glenda Gray. PP/COS cohorts were managed by Katherine Thomas, Connie Celum, Nelly Mugo, Andrew Mujugira, Jairam Lingappa. Writing and editing of the original manuscript draft was done by Tosin E. Omole, Lyle R. McKinnon. All authors have reviewed, edited, and approved the manuscript.

## Acknowledgements

We would like to acknowledge all participants who donated their specimens for this research.

## Supporting information

**S1 Fig. Representative flow cytometry plots illustrating the gating strategy for IL-17^+^ CD4^+^ T cells and IL-17^+^ cytokine co-expressing subsets.**

**S2 Fig. Representative flow cytometry plots illustrating the gating strategy for non-IL-17-gated CD4^+^ T cell cytokine subsets.**

**S3 Fig. Correlation between mean CD4/CD8 ratio within the first 180 days post-infection and pre-HIV IL-17^+^ CD4^+^ T cell cytokine co-expressing subsets in HVTN 503.** Panels show correlations for the following IL-17^+^ **subsets: (A)** TNF-α^+^, **(B)** TNF-α^-^, **(C)** IFN-γ^+^, **(D)** IFN-γ^-^, **(E)** GM-CSF^+^, **(F)** GM-CSF^-^, **(G)** IL22^+^, **(H)** IL22^-^. CD4/CD8 ratio was calculated from absolute CD4 and CD8 counts measured within the initial 180 days post-infection. CD4/CD8 ratios were calculated from absolute CD4 and CD8 counts measured within the first 180 days post-infection (n = 35). Measurements obtained after ART initiation or beyond 1-year post-infection were excluded. Correlations were assessed using Spearman rank correlation (R), with linear regression lines shown for visualization. Two-tailed *p* values are shown; statistical significance was defined as *p* < 0.05.

**S4 Fig. Correlation between mean CD4/CD8 ratio beyond the first 180 days post-infection and pre-HIV IL-17^+^ CD4^+^ T cell cytokine co-expressing subsets in HVTN 503.** Panels show correlations for the following IL-17^+^ subsets: (A) TNF-α^+^, (B) TNF-α^-^, (C) IFN-γ^+^, (D) IFN-γ^-^, (E) GM-CSF^+^, (F) GM-CSF^-^, (G) IL22^+^, (H) IL22^-^. CD4/CD8 ratio was calculated from absolute CD4 and CD8 counts measured after the initial 180 days post-infection. CD4/CD8 ratios were calculated from absolute CD4 and CD8 counts measured within the first 180 days post-infection (n = 32). Measurements obtained after ART initiation or beyond 1-year post-infection were excluded. Correlations were assessed using Spearman rank correlation (R), with linear regression lines shown for visualization. Two-tailed *p* values are shown; statistical significance was defined as *p* < 0.05.

**S5 Fig. Correlation between mean CD4/CD8 ratio and pre-HIV non-IL-17-gated CD4^+^ T cell cytokine subsets in HVTN 503.** Panels show correlations with mean CD4/CD8 ratio measured: (A – D) within the first 180 days post-infection (n = 35), (E – H) after the first 180 days post-infection subset (n = 32). CD4/CD8 ratios were calculated from absolute CD4 and CD8 counts measured within and after the first 180 days post-infection. Measurements obtained after ART initiation or beyond 1-year post-infection were excluded. Correlations were assessed using Spearman rank correlation (R), with linear regression lines shown for visualization. Two-tailed *p* values are shown; statistical significance was defined as p < 0.05.

**S6 Fig. Sex-stratified correlation between mean CD4/CD8 ratio within the first 180 days post-infection and pre-HIV IL-17^+^ CD4^+^ T cell cytokine co-expressing subsets in HVTN 503.** CD4/CD8 ratios were calculated from absolute CD4 and CD8 counts measured within the first 180 days post-infection (n = 35). Measurements obtained after ART initiation or beyond 1-year post-infection were excluded. Correlations were assessed using Spearman rank correlation (R), with linear regression lines shown for visualization. Two-tailed *p* values are shown; statistical significance was defined as *p* < 0.05.

**S7 Fig. Sex-stratified correlation between mean CD4/CD8 ratio beyond the first 180 days post-infection and pre-HIV IL-17^+^ CD4^+^ T cell cytokine co-expressing subsets in HVTN 503.** CD4/CD8 ratios were calculated from absolute CD4 and CD8 counts measured after the initial 180 days post-infection (n = 32). Measurements obtained after ART initiation or beyond 1-year post-infection were excluded. Correlations were assessed using Spearman rank correlation (R), with linear regression lines shown for visualization. Two-tailed *p* values are shown; statistical significance was defined as *p* < 0.05.

**S8 Fig. Treatment arm-stratified correlation between mean CD4/CD8 ratio within the first 180 days post-infection and pre-HIV IL-17^+^ CD4^+^ T cell cytokine co-expressing subsets in HVTN 503.** CD4/CD8 ratios were calculated from absolute CD4 and CD8 counts measured within the first 180 days post-infection (n = 35). Measurements obtained after ART initiation or beyond 1-year post-infection were excluded. Correlations were assessed using Spearman rank correlation (R), with linear regression lines shown for visualization. Two-tailed *p* values are shown; statistical significance was defined as *p* < 0.05.

**S9 Fig. Treatment arm-stratified correlation between mean CD4/CD8 ratio beyond the first 180 days post-infection and pre-HIV IL-17^+^ CD4^+^ T cell cytokine co-expressing subsets in HVTN 503.** CD4/CD8 ratios were calculated from absolute CD4 and CD8 counts measured after the initial 180 days post-infection (n = 32). Measurements obtained after ART initiation or beyond 1-year post-infection were excluded. Correlations were assessed using Spearman rank correlation (R), with linear regression lines shown for visualization. Two-tailed *p* values are shown; statistical significance was defined as *p* < 0.05.

**S10 Fig. Correlation between peak viral load and pre-HIV IL-17^+^ CD4^+^ T cell cytokine co-expressing subsets.** Data from HVTN 503 are shown in red (n = 35) and PP/COS in blue (n = 32). Peak viral load was defined as the highest viral load measured within the first 180 days post-infection. Viral load measurements obtained after ART initiation or beyond 1-year post-infection were excluded. Correlations were assessed using Spearman rank correlation (R), with linear regression lines shown for visualization. Two-tailed *p* values are shown; statistical significance was defined as *p* < 0.05.

**S11 Fig. Correlation between set-point viral load and pre-HIV IL-17^+^ CD4^+^ T cell cytokine co-expressing subsets.** Data from HVTN 503 are shown in red (n = 32) and PP/COS in blue (n = 18). Set-point viral load was defined as the mean viral load measured after 180 days post-infection. Viral load measurements obtained after ART initiation or beyond 1-year post-infection were excluded. Correlations were assessed using Spearman rank correlation (R), with linear regression lines shown for visualization. Two-tailed *p* values are shown; statistical significance was defined as *p* < 0.05.

**S1 Table. Association between pre-HIV IL-17^+^ CD4^+^ T cells and CD4 decline below 500 cells/mm^3^, adjusted for viral load (HVTN 503).** Hazard ratios were estimated using Cox proportional hazards model. The model was adjusted only for peak viral load. Two-tailed *p* values are shown; statistical significance was defined as *p* < 0.05. Abbreviations: aHR = Adjusted Hazard Ratio; CI = Confidence Interval; HVTN = HIV Vaccine Trials Network.

**S2 Table. Association between pre-HIV IL-17^+^ CD4^+^ T cells and CD4 decline below 500 cells/mm^3^, adjusted for covariates excluding viral load (HVTN 503).** Hazard ratios were estimated using Cox proportional hazards model. The model was adjusted for sex, adenovirus type 5 (Ad5) titer, herpes simplex virus-2 (HSV-2) status, and age. Two-tailed *p* values are shown; statistical significance was defined as *p* < 0.05. Abbreviations: aHR = Adjusted Hazard Ratio; CI = Confidence Interval; HVTN = HIV Vaccine Trials Network.

**S3 Table. Association between pre-HIV non-IL-17-gated CD4^+^ T cell cytokine subsets and CD4 decline below 500 cells/mm^3^.** Frequency of each phenotype below the median value was used as the reference category. Hazard ratios were estimated using unadjusted Cox proportional hazards model. Two-tailed *p* values are shown; statistical significance was defined as *p* < 0.05. Abbreviations: HR = Hazard Ratio; CI = Confidence Interval; HVTN = HIV Vaccine Trials Network; PP/COS = Partners PrEP/Couples Observational Study.

**S4 Table. Intracellular cytokine staining panel for ex vivo Th17 CD4^+^ T cell immunophenotyping in PBMC samples**

**S5 Table. General reagents used for sample processing and data acquisition**

**S6 Table. Stimulation reagents used for ex vivo Th17 immunophenotyping in PBMC samples**

## References

1. Sivro A, Schuetz A, Sheward D, Joag V, Yegorov S, Liebenberg LJ, et al. Integrin α4β7 expression on peripheral blood CD4+ T cells predicts HIV acquisition and disease progression outcomes. Sci Transl Med. 2018;10: eaam6354. doi:10.1126/scitranslmed.aam6354

2. Hartigan-O’Connor DJ, Abel K, Van Rompay KKA, Kanwar B, McCune JM. SIV replication in the infected rhesus macaque is limited by the size of the preexisting T H17 cell compartment. Sci Transl Med. 2012;4: 136ra69. doi:10.1126/scitranslmed.3003941

3. Koning FA, Otto SA, Hazenberg MD, Dekker L, Prins M, Miedema F, et al. Low-Level CD4 T Cell Activation Is Associated with Low Susceptibility to HIV-1 Infection 1. 2005;175: 6117–6122.

4. Ngcobo S, Molatlhegi RP, Osman F, Ngcapu S, Samsunder N, Garrett NJ, et al. Pre-infection plasma cytokines and chemokines as predictors of HIV disease progression. Sci Rep. 2022;12: 2437. doi:10.1038/s41598-022-06532-w

5. Omole TE, Nguyen HM, Marcinow A, Oo MM, Jahan N, Ssemaganda A, et al. Pre–Human Immunodeficiency Virus (HIV) α4β7hi CD4+ T Cells and HIV Risk Among Heterosexual Individuals in Africa. Journal of Infectious Diseases. 2025;231: e770–e780. doi:10.1093/infdis/jiae638

6. Omole TE, McKinnon LR. Integrin α4β7 as a predictor of HIV acquisition: one thread in a complex tapestry. Journal of Clinical Investigation. American Society for Clinical Investigation; 2025. doi:10.1172/JCI195258

7. Roberts L, Passmore JAS, Williamson C, Little F, Bebell LM, Mlisana K, et al. Plasma cytokine levels during acute HIV-1 infection predict HIV disease progression. AIDS. 2010;24: 819–831. doi:10.1097/QAD.0b013e3283367836

8. Klatt NR, Brenchley JM. Th17 cell dynamics in HIV infection. Curr Opin HIV AIDS. 2010;5: 135–140. doi:10.1097/COH.0b013e3283364846

9. Paiva IA, Badolato-Corrêa J, Familiar-Macedo D, De-Oliveira-pinto LM. Th17 cells in viral infections—friend or foe? Cells. 2021;10: 1159. doi:10.3390/cells10051159

10. Rodriguez-Garcia M, Barr FD, Crist SG, Fahey J V., Wira CR. Phenotype and susceptibility to HIV infection of CD4+ Th17 cells in the human female reproductive tract. Mucosal Immunol. 2014;7: 1375–1385. doi:10.1038/mi.2014.26

11. Muranski P, Restifo NP. Essentials of Th17 cell commitment and plasticity. Blood. 2013;121: 2402–2414. doi:10.1182/blood

12. Wacleche VS, Landay A, Routy JP, Ancuta P. The Th17 lineage: From barrier surfaces homeostasis to autoimmunity, cancer, and HIV-1 pathogenesis. Viruses. 2017;9: 303. doi:10.3390/v9100303

13. Ancuta P, Monteiro P, Sekaly RP. Th17 lineage commitment and HIV-1 pathogenesis. Curr Opin HIV AIDS. 2010;5: 158–165. doi:10.1097/COH.0b013e3283364733

14. Mckinnon LR, Nyanga B, Kim CJ, Izulla P, Kwatampora J, Kimani M, et al. Early HIV-1 Infection Is Associated with Reduced Frequencies of Cervical Th17 Cells. Journal of Acquired Immune Deficiency Syndrome. 2015;68: 6–12.

15. Maric D, Grimm WA, Greco N, Mcraven MD, Fought AJ, Veazey RS, et al. Th17 T Cells and Immature Dendritic Cells Are the Preferential Initial Targets after Rectal Challenge with a Simian Immunodeficiency Virus-Based Replication-Defective Dual-Reporter Vector. J Virol. 2021;95: e00707–21.

16. Stieh DJ, Matias E, Marx PA, Veazey RS, Correspondence TJH. Th17 Cells Are Preferentially Infected Very Early after Vaginal Transmission of SIV in Macaques. Cell Host Microbe. 2016;19: 529–540. doi:10.1016/j.chom.2016.03.005

17. Schuetz A, Deleage C, Sereti I, Rerknimitr R, Phanuphak N, Phuang-Ngern Y, et al. Initiation of ART during Early Acute HIV Infection Preserves Mucosal Th17 Function and Reverses HIV-Related Immune Activation. PLoS Pathog. 2014;10: e1004543. doi:10.1371/journal.ppat.1004543

18. Loiseau C, Requena M, Mavigner M, Cazabat M, Carrere N, Suc B, et al. CCR6-regulatory T cells blunt the restoration of gut Th17 cells along the CCR6-CCL20 axis in treated HIV-1-infected individuals. Mucosal Immunol. 2016;9: 1137–1150. doi:10.1038/mi.2016.7

19. Renault C, Veyrenche N, Mennechet F, Bedin AS, Routy JP, Van de Perre P, et al. Th17 CD4+ T-Cell as a Preferential Target for HIV Reservoirs. Front Immunol. 2022;13: 822576. doi:10.3389/fimmu.2022.822576

20. Zayas JP, Mamede JI. HIV Infection and Spread between Th17 Cells. Viruses. 2022;14: 404. doi:10.3390/v14020404

21. Fernandes JR, Berthoud TK, Kumar A, Angel JB. IL-23 signaling in Th17 cells is inhibited by HIV infection and is not restored by HAART: Implications for persistent immune activation. PLoS One. 2017;12: e0186823. doi:10.1371/journal.pone.0186823

22. Wiche-Salinas T. R., Zhang Y, Sarnello D, Zhyvoloup A, Marchand LR, Fert A, et al. Th17 cell master transcription factor RORC2 regulates HIV-1 gene expression and viral outgrowth. PNAS. 2021;118: e2105927118. doi:10.1073/pnas.2105927118/-/DCSupplemental

23. Gray GE, Moodie Z, Metch B, Gilbert PB, Bekker LG, Churchyard G, et al. The phase 2b HVTN 503/Phambili study test-of-concept HIV vaccine study, investigating a recombinant adenovirus type 5 HIV gag/pol/nef vaccine in South Africa: unblinded, long-term follow-up. Lancet Infect Dis. 2014;14: 388–396. doi:10.1016/S1473-3099(14)70020-9

24. Machmach K, N’guessan KF, Farmer R, Godbole S, Kim D, McCormick L, et al. NK cell activation and CD4 T cell α4β7 expression are associated with susceptibility to HIV-1. Journal of Clinical Investigation. 2025. doi:10.1172/JCI187992

25. El Hed A, Khaitan A, Kozhaya L, Manel N, Daskalakis D, Borkowsky W, et al. Susceptibility of human Th17 cells to human immunodeficiency virus and their perturbation during infection. Journal of Infectious Diseases. 2010;201: 843–854. doi:10.1086/651021

26. Guillot-Delost M, Le Gouvello S, Mesel-Lemoine M, Cheraï M, Baillou C, Simon A, et al. Human CD90 Identifies Th17/Tc17 T Cell Subsets That Are Depleted in HIV-Infected Patients. The Journal of Immunology. 2012;188: 981–991. doi:10.4049/jimmunol.1101592

27. Anastos K, Gange SJ, Lau B, Weiser B, Detels R, Giorgi J V., et al. Association of Race and Gender With HIV-1 RNA Levels and Immunologic Progression. Journal of Acquired Immune Deficiency Syndrome. 2000;24: 218–226.

28. D’Ettorre G, Borrazzo C, Pinacchio C, Santinelli L, Cavallari EN, Statzu M, et al. Increased IL-17 and/or IFN-γ producing T-cell subsets in gut mucosa of long-term-treated HIV-1-infected women. AIDS. 2019;33: 627–636. doi:10.1097/QAD.0000000000002122

29. Cohn J, Ake J, Moorhouse M, Godfrey C. Sex Differences in the Treatment of HIV. Curr HIV/AIDS Rep. 2020;17: 373–384. doi:10.1007/s11904-020-00499-x

30. Mosha F, Muchunguzi V, Matee M, Sangeda RZ, Vercauteren J, Nsubuga P, et al. Gender differences in HIV disease progression and treatment outcomes among HIV patients one year after starting antiretroviral treatment (ART) in Dar es Salaam, Tanzania. BMC Public Health. 2013;13: 38. doi:10.1186/1471-2458-13-38

31. Fuseini H, Yung JA, Cephus JY, Zhang J, Goleniewska K, Polosukhin V V, et al. Testosterone Decreases House Dust Mite–Induced Type 2 and IL-17A–Mediated Airway Inflammation. The Journal of Immunology. 2018;201: 1843–1854. doi:10.4049/jimmunol.1800293

32. Dodd KC, Menon M. Sex bias in lymphocytes: Implications for autoimmune diseases. Frontiers in Immunology. Frontiers Media S.A.; 2022. doi:10.3389/fimmu.2022.945762

33. Ngo ST, Steyn FJ, McCombe PA. Gender differences in autoimmune disease. Frontiers in Neuroendocrinology. Academic Press Inc.; 2014. pp. 347–369. doi:10.1016/j.yfrne.2014.04.004

34. Mani NL, Weinberg SE. Sex, cells, and metabolism: Androgens temper Th17-mediated immunity. Journal of Clinical Investigation. 2024;134: e186520. doi:10.1172/JCI177242

35. Chowdhury NU, Cephus JY, Pilier EH, Wolf MM, Madden MZ, Kuehnle SN, et al. Androgen signaling restricts glutaminolysis to drive sex-specific Th17 metabolism in allergic airway inflammation. Journal of Clinical Investigation. American Society for Clinical Investigation; 2024. p. e177242. doi:10.1172/JCI186520

36. Trigunaite A, Dimo J, Jørgensen TN. Suppressive effects of androgens on the immune system. Cell Immunol. 2015;294: 87–94. doi:10.1016/J.CELLIMM.2015.02.004

37. Klein SL, Flanagan KL. Sex differences in immune responses. Nat Rev Immunol. 2016;16: 626–638. doi:10.1038/NRI.2016.90;SUBJMETA

38. Wilkinson NM, Chen HC, Lechner MG, Su MA. Sex Differences in Immunity. Annual Review of Immunology. Annual Reviews Inc.; 2022. pp. 75–94. doi:10.1146/annurev-immunol-101320-125133

39. Baeten JM, Donnell D, Ndase P, Mugo NR, Campbell JD, Wangisi J, et al. Antiretroviral Prophylaxis for HIV Prevention in Heterosexual Men and Women. New England Journal of Medicine. 2012;367: 399–410. doi:10.1056/nejmoa1108524

40. Hopkins KL, Laher F, Otwombe K, Churchyard G, Bekker LG, DeRosa S, et al. Predictors of HVTN 503 MRK-AD5 HIV-1 gag/pol/nef vaccine induced immune responses. PLoS One. 2014;9: e103446. doi:10.1371/journal.pone.0103446

41. Mujugira A, Baeten JM, Donnell D, Ndase P, Mugo NR, Barnes L, et al. Characteristics of HIV-1 serodiscordant couples enrolled in a clinical trial of antiretroviral pre-exposure prophylaxis for HIV-1 prevention. PLoS One. 2011;6: e25828. doi:10.1371/journal.pone.0025828

42. Murnane PM, Celum C, Mugo N, Campbell JD, Donnell D, Bukusi E, et al. Efficacy of preexposure prophylaxis for HIV-1 prevention among high-risk heterosexuals: Subgroup analyses from a randomized trial. AIDS. 2013;27: 2155–2160. doi:10.1097/QAD.0b013e3283629037

